# μ-Opioid receptor transcriptional variants in the murine forebrain and spinal cord

**DOI:** 10.1101/2024.03.20.585849

**Authors:** Magdalena Chrószcz, Jacek Hajto, Klaudia Misiołek, Łukasz Szumiec, Magdalena Ziemiańska, Anna Radlicka-Borysewska, Małgorzata Borczyk, Mateusz Zięba, Sławomir Gołda, Marcin Siwiec, Barbara Ziółkowska, Marcin Piechota, Michał Korostyński, Jan Rodriguez Parkitna

**Author notes:** Corresponding Address: Jan Rodriguez Parkitna, Department of Molecular Neuropharmacology, Maj Institute of Pharmacology, Polish Academy of Sciences Smętna 12, 31-343, Krakow, Poland.

## Abstract

**Background:** *Oprm1*, the gene encoding the μ-opioid receptor, has multiple reported transcripts, with a variable 3 region and many alternative sequences encoding the C-terminus of the protein. The functional implications of this variability remain mostly unexplored, though a recurring notion is that it could be exploited by developing selective ligands with improved clinical profiles. Here, we comprehensively examined *Oprm1* transcriptional variants in the murine central nervous system.

**Methods:** RNA-seq transcription analyses were performed based on Oxford Nanopore Sequencing (ONS) and 10x Genomics Visium spatial transcriptomics data. The spatial distribution of *Oprm1* exons was evaluated via RNAscope *in situ* hybridization. Tissue and cell-type specificity was assessed based on reanalysis single-cell RNAseq databases.

**Results:** We detected a mismatch between transcripts annotated in GRCm38/mm10 and RNA-seq results. Sequencing data indicated that the primary *Oprm1* transcript has a 3 terminus located on chr10:6,860,027, which is ~9.5 kilobases downstream of the longest annotated exon 4 end. Long-read sequencing confirmed that the final *Oprm1* exon included a 10.2 kilobase long 3 untranslated region. The presence of the long variant was unambiguously confirmed using RNAscope *in situ* hybridization. The long variant was observed in the thalamus, striatum, cortex and spinal cord. Expression of additional variants of the *Oprm1* gene was close to the detection limit. Reanalysis of single-cell sequencing data confirmed these observations and indicated that *Oprm1* was expressed mainly in parvalbumin-, somatostatin- and VIP-positive cells.

**Conclusion:** The primary transcript of the *Oprm1* mouse gene is a variant with a long 3 untranslated region.

**Author Summary:** Opioids are essential for the management of pain and have multiple other medical indications; however, their addictive properties and widespread misuse have led to a severe modern health crisis. Accordingly, there has been a major effort to develop novel compounds that retain clinical effectiveness while diminishing their addictive potential and other adverse effects. One of the potential avenues for safer opioid drugs is developing compounds that are selective for a specific group of the main targets of opioid medications—the μ-opioid receptors. Multiple variants the μ-opioid receptor have been reported, encoded by different transcripts of the *Oprm1* gene. Here, we used RNA transcript sequencing and *in situ* hybridization with probes to detect different parts of *Oprm1* transcripts to validate the existence of various reported isoforms. Our main finding is that the primary transcript of the receptor is much longer than the current reference sequences annotated in the mouse genome and has an over 10,000-base-long noncoding sequence at the 3 terminus. Several other types of transcripts are also expressed; however, they represent approximately 15% or less of the total transcript content in each of the examined brain regions. In the context of future research on opioid drugs, these results indicate that it is unlikely that different subpopulations of receptors could be targeted.

## Introduction

The murine *Oprm1* gene encodes the μ-opioid receptor, which belongs to a rhodopsin-like G-protein coupled receptor (GPCR) family and is primarily bound to a G_i/o_ heterotrimer. The receptor was first cloned in rats (Chen et al., 1993) and subsequently cloned in humans and mice (Min et al., 1994; Wang et al., 1994). Cloning of the complete receptor sequence led to the generation of gene-knockout models and the seminal observation that the analgesic effects of morphine are mediated by the μ-opioid receptor (Loh et al., 1998; Matthes et al., 1996). The essential role of the μ-opioid receptor in analgesia remains the driver for extensive research on μ-opioid receptor signaling and the development of novel synthetic opioids. Nevertheless, it should be noted that μ-opioid receptors also play major roles in the control of the reward system of the brain, the modulation of affective states, the stress response, and a wide array of peripheral functions, e.g., the regulation of enteric motility or the immune response (Berrendero et al., 2002; Eisenstein, 2019; Filliol et al., 2000; Le Merrer et al., 2009; Moles, 2004; Wood & Galligan, 2004).

The murine *Oprm1* gene is located on chromosome 10 (10qA1) and is ubiquitously expressed across tissues. The highest transcript abundance was observed in the central nervous system, particularly in the thalamus, hypothalamus, basal ganglia, midbrain and superficial layers of the dorsal horn of the spinal cord (Delfs et al., 1994; Mansour, Fox, et al., 1995; Märtin et al., 2019; Thompson et al., 1993). On the periphery, robust expression of the receptor was observed mainly in the testes (Estomba et al., 2016), with some studies suggesting its expression in activated immune cells as well (Machelska & Celik, 2020; Zhang et al., 2012). The μ-opioid receptor appears to have low selectivity for endogenous opioid peptides (Mansour, Hoversten, et al., 1995) and was reported to show heterogeneous responses to synthetic opioids (Chang et al., 1998; Pasternak, 2012; Paul et al., 1989; Rivero et al., 2012). These findings, along with incomplete cross-tolerance after chronic exposure to selective μ-opioid agonists, led to ongoing speculation on the existence of functionally different receptor isoforms (Goldstein & James, 1984; Liu et al., 2021; Pasternak, 2001).

This idea aligns with findings that have shown the existence of alternative *Oprm1* transcripts. In the past 3 decades, more than 90 alternative *Oprm1* transcript variants have been described in mice (*Mus musculus*) with putative alternative starts (exons 1 & 11), a conserved sequence encoding transmembrane helices (exons 2 & 3), and highly variable 3 termini encoding part of the protein C-terminus and the 3 UTR (Doyle, Rebecca Sheng, et al., 2007; Doyle, Sheng, et al., 2007; Pan et al., 1999, 2000, 2001, 2005). A summary of *Oprm1* transcripts present in the GenBank and ENSEMBL databases is shown in **Figure 1**. The majority of reported protein-coding sequences differ in their C-terminal sequences (Pasternak & Pan, 2013), although the expression of truncated but functional receptors comprising only 6 transmembrane helices has also been reported (Majumdar et al., 2011; Xu et al., 2013). Furthermore, it was reported that the *Oprm1* transcript in the brain contains a 10 kb long 3 terminal sequence based on Northern blot and sequencing of PCR fragments (Ide et al., 2005; Ikeda et al., 2001; Wu et al., 2005). It should be noted, however, that the isoform with a long 3 end was not further characterized, nor included in current mouse genomic databases (Ensembl release 108 and GenBank Release 253).

**Figure 1.**
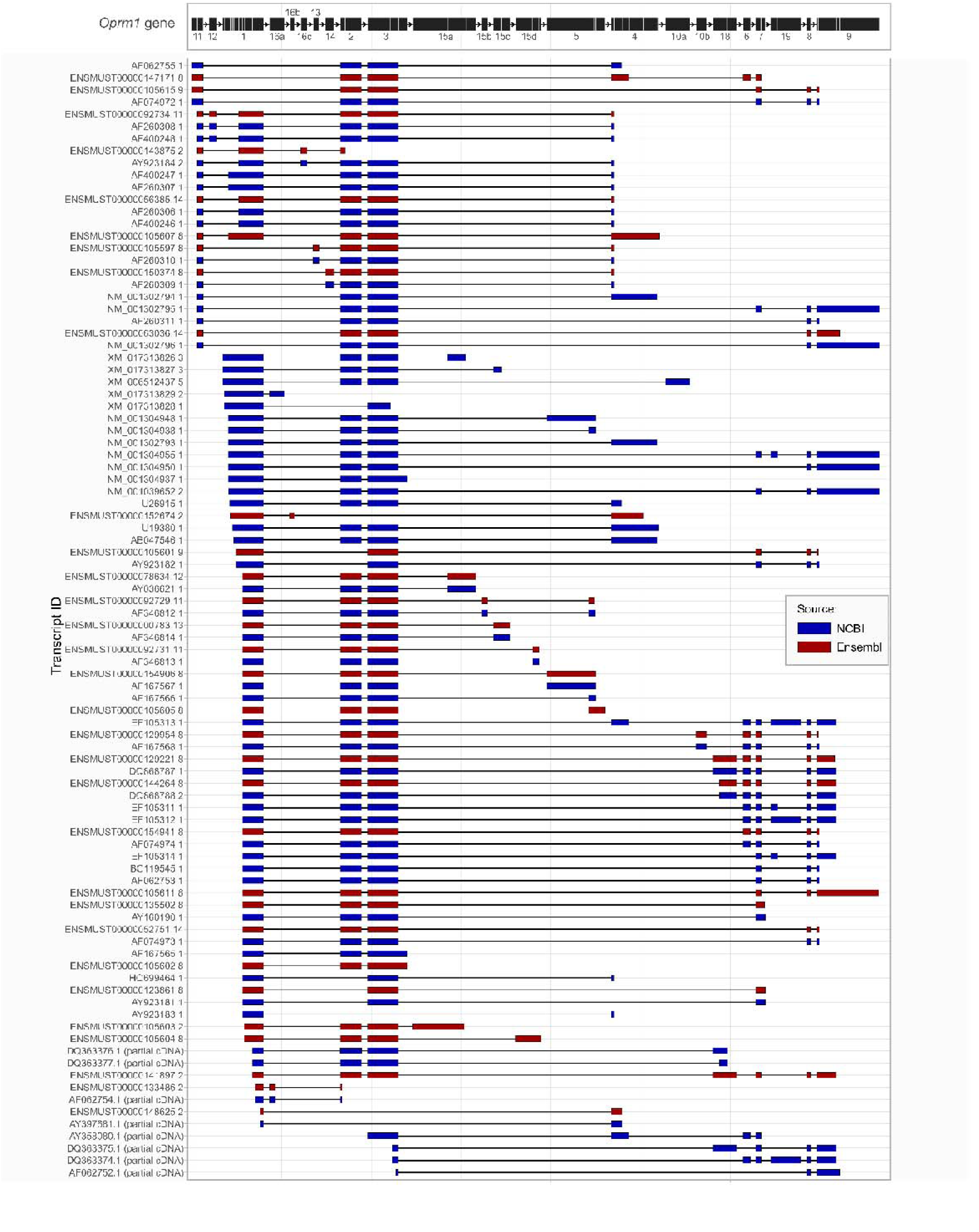
Schematic representation of *Oprm1* gene transcripts reported in the Ensembl and GenBank databases. The transcripts shown are based on Ensembl release 108 and GenBank release 253. The diagram at the top of the image shows a summary of reported exons. Identifiers for each transcript are shown on the left, boxes represent exons, and colors correspond to the source of the transcript annotation.

The variability of the C-terminus is particularly interesting, as this domain likely remains in proximity to the G protein complex and could play a role in the bias toward activation of the β-arrestin pathway (Kliewer et al., 2019; Narayan et al., 2021). However, it should be noted that the position of the C-terminus is not included in the available crystal structures (Koehl et al., 2018; Manglik et al., 2012); thus, its role in the interaction with downstream signaling pathways remains speculative. The putative presence of isoforms differing in their ligand affinity and ability to transactivate G-protein or β-arrestin pathways and expressed in different cell types offers a plausible explanation for some of the differences in the action of opioid drugs and a theoretical possibility for the design of novel opioids with reduced adverse effects.

Here, we comprehensively examined the transcription of *Oprm1* mRNA in the mouse brain. In line with some previous reports, we found a single primary transcript of the *Oprm1* gene, which includes an ultralong 3 UTR with a short C-terminus that is highly conserved between mice and humans. Other transcripts with alternate 3 ends were also detected; however, we estimated that their abundance was considerably lower than that of the main transcript and accounted for 5–20% of the overall *Oprm1* expression. Reanalysis of Allen Atlas single-cell sequencing data confirmed the presence of the novel transcript in selected types of cortical neurons.

## Results

### Mouse Oprm1 transcripts

*Oprm1* transcription was previously extensively characterized using *in situ* hybridization with radioactive cDNA probes and PCR (Doyle, Sheng, et al., 2007; Ikeda et al., 2001; Kitchen et al., 1997; Mansour, Fox, et al., 1995; Matthes et al., 1996; Moskowitz & Goodman, 1984; Pan et al., 1999; Sharif & Hughes, 1989). Here, we first assessed to what extent the previously reported data aligned with available single-cell RNA-seq datasets (**Figure 2**). Reanalysis of the Tabula Muris dataset revealed that reads aligned to *Oprm1* exons were found primarily in neurons (**Figure 2**) and in several immune and progenitor cell types, including granulocytes, granulopoietic cells, natural killer cells, professional antigen-presenting cells, *Slamf1*-negative multipotent progenitors, and megakaryocyte-erythroid progenitors. However, the Tabula Muris dataset does not distinguish between DNA strands, and in the case of non-neuronal cells, the reads were aligned primarily in the region corresponding to exons 6 to 10, which overlap with the *Ipcef1* gene (encoded on the opposite strand from *Oprm1*). Altogether, most-conserved *Oprm1* exons 1 to 3 were unambiguously present only in neurons, with a mean coverage for these exons equal to 0.4192 per cell (**Figure 2**). Low expression of all these exons was also detected in endothelial cells, but the values were close to the detection limit (mean coverage per cell of 0.0005). It is important to note that some types of non-neuronal cells known to express high levels of μ-opioid receptors, e.g., Sertoli cells or spermatids, were not present in the Tabula Muris dataset at the time the analyses were performed. In summary, the analysis of the dataset confirmed that *Oprm1* is expressed primarily in neurons, with strong evidence for transcripts including exons 1, 2 and 3 but no clear evidence for the presence of 3 terminal regions.

**Figure 2.**
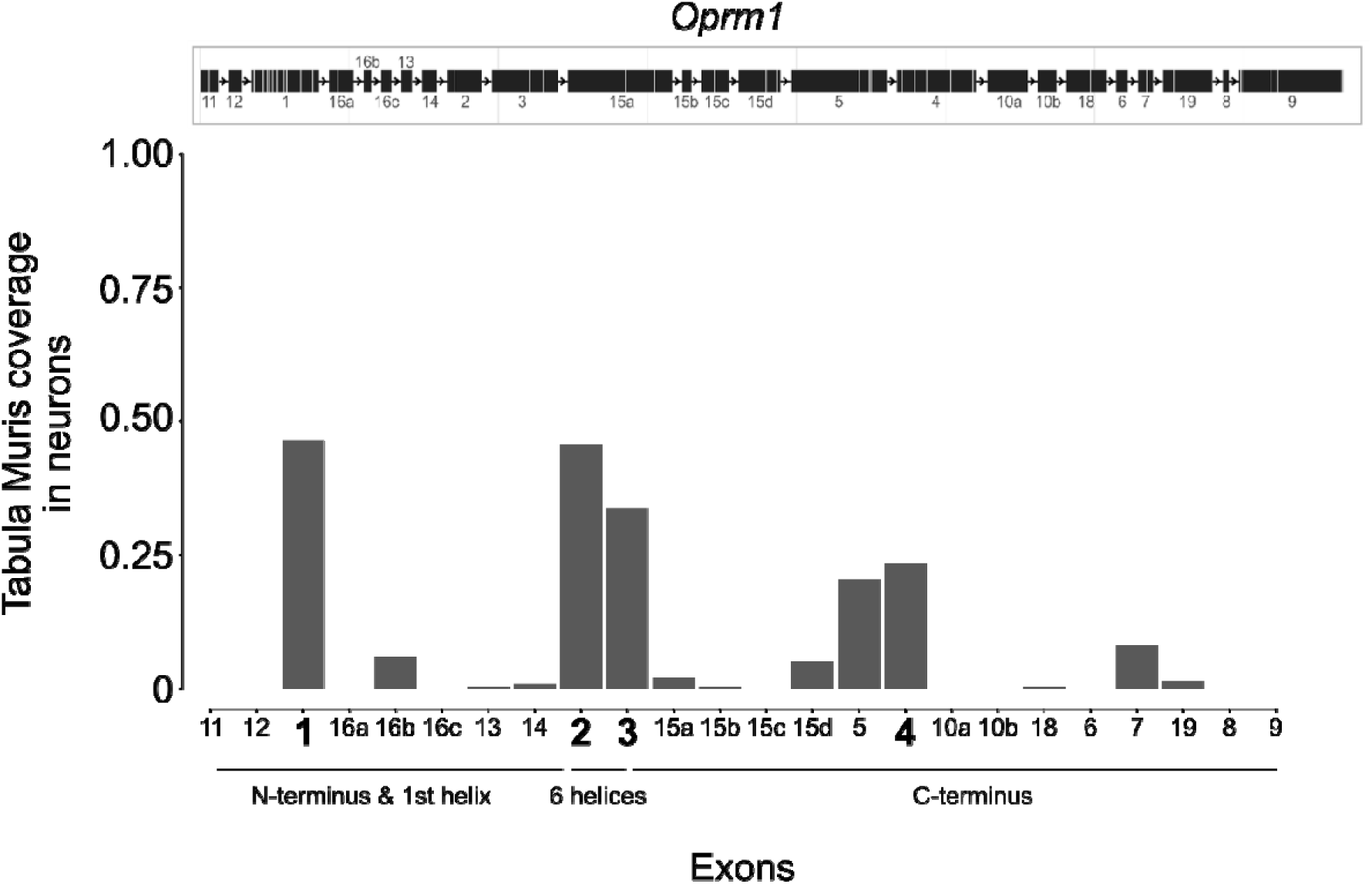
Alignment of sequence reads from the Tabula Muris dataset in neuronal cells to putative *Oprm1* exons. The diagram at the top represents reported exons and their order. The bar graph shows relative coverage in regions corresponding to specific exons. Due to differences in exon length for different reported transcripts, the most conserved exon ranges were included in the analysis, as depicted in pink on the diagram. Bars corresponding to the sequence of exons 1 to 4 present in the reported mouse sequence are shown in black.

### The primary Oprm1 transcript in the mouse forebrain includes a 10-kbp long 30 UTR

Next, to directly map *Oprm1* transcript variants in the brain, we performed long-read RNA sequencing using the Oxford Nanopore platform. RNA samples for sequencing were prepared from male and female coronal mouse brain slices 0,74-1.34 mm anterior to bregma (which included the striatum and the medial prefrontal cortex), and the thalamus was extracted with needles from slices –0.94 to –1.82 mm posterior to bregma. RNA was purified, reverse-transcribed into cDNA, and sequenced using the PromethION platform, and the obtained reads were aligned to the GRCm38/mm10 mouse genome. The alignment of the sequenced reads on chromosome 10qA1 is shown in **Figures 3** and **S1**. Reads cover *Oprm1* exons 1 through 4 and extend over an ~10 kb sequence beyond the annotated 3 end of exon 4. The read coverage of this region was uneven, but the analysis suggested the presence of a single long exon, both in the prefrontal cortex/striatum and the thalamus. A few reads were aligned in the region corresponding to *Oprm1* exons 7 to 9 (**Figure S1**); however, they constituted less than 5% of the reads aligned to exons 2 and 3. Thus, long-read sequencing revealed that exon 4 of *Oprm1* extends an additional 10 kb in the 3 direction and that the majority of *Oprm1* transcripts include long exon 4. No evidence of alternative transcription start sites was observed.

**Figure 3.**
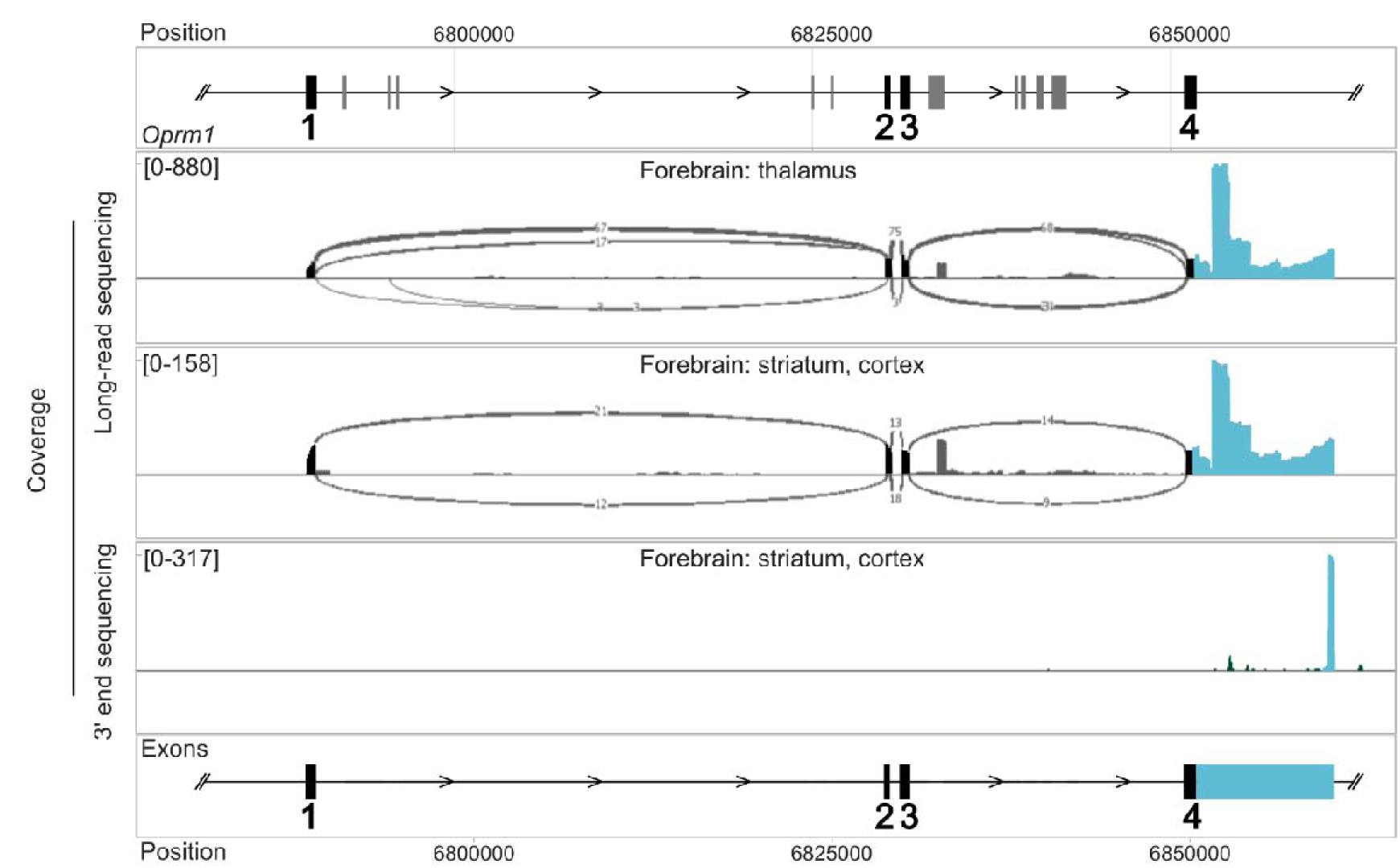
RNA sequencing analysis of *Oprm1* transcripts in the mouse forebrain. The diagram at the top shows the positions of putative exons in the 5’ region of the gene. The position corresponds to chromosome 10qA1. The first two lanes show “Sashimi” plots of long-read sequencing analysis of brain regions, including the cortex and striatum (the first lane) and the thalamus (the second lane). The third lane shows sequencing results based on spatial RNA sequencing of 3’ termini in a frontal brain coronal section (spatial results from the whole brain merged). The diagram at the bottom shows the transcripts reconstructed based on the sequencing results.

To map the 3 termini of *Oprm1* transcripts, we used RNA-seq data from spatial transcriptomics analyses performed on coronal mouse brain slices that included the striatum and prefrontal cortex. The data acquisition utilized the Visium method and thus consisted of a large number of stranded reads of the 3 ends of transcripts. The results are summarized in **Figure 3** and **Figure S1**. The primary putative 3 terminus was located at position 6,859,750–6,860,000 bp (317 reads) and aligned with the end of the extended 3 exon identified via long-read sequencing (**Figure 3**). Additional putative 3 termini were also identified at positions 6,852,700–6,852,900 bp (40 reads), 6,985,900–6,986,200 bp (59 reads), and 7,038,750–7,038,830 bp (762 reads). The first two were located in proximity to the reported exon 19, with the exon end at 6,985,900–6,986,200. The 7,038,750–7,038,830 region is in proximity to exon 9; however, these reads were aligned to the opposite strand. In summary, the alignment of the 3-end sequencing data indicated that exon 4 is significantly longer than previously annotated, encompassing a 10 kbp untranslated region. Again, the presence of alternative 3-end variants was confirmed, but they constituted a relatively minor fraction (16,5% assuming no false-positive results) of the transcripts detected.

This novel *Oprm1* variant is consistent with previous reports from Ikeda and colleagues (Ide et al., 2005), which showed that long exon 4 was homologous to the NM000914 reference transcript of human *OPRM1*. A comparison of the human and murine transcripts is shown in **Figure 4**. There is a clear similarity in the sequence corresponding to exons 1, 2, and 3 (until base position ~4000 in the mouse transcript), which also extends to the 51 region of the long exon 4. Additionally, the 3 region of long exon 4 (starting approximately 7000 bases from the 3 end of the mouse transcript) appears to be conserved. The extended 3’ final exon emerges as a distinctive characteristic of *Oprm1*, settling it apart from other opioid receptor genes. Analysis of the long sequence reads of its paralogs, i.e., *Oprd1*, *Oprk1,* and *Oprl1,* confirmed that the final exons of these genes were shorter, with 3 termini and lengths ranging between 1,600 bp and 4,200 bp.

**Figure 4.**
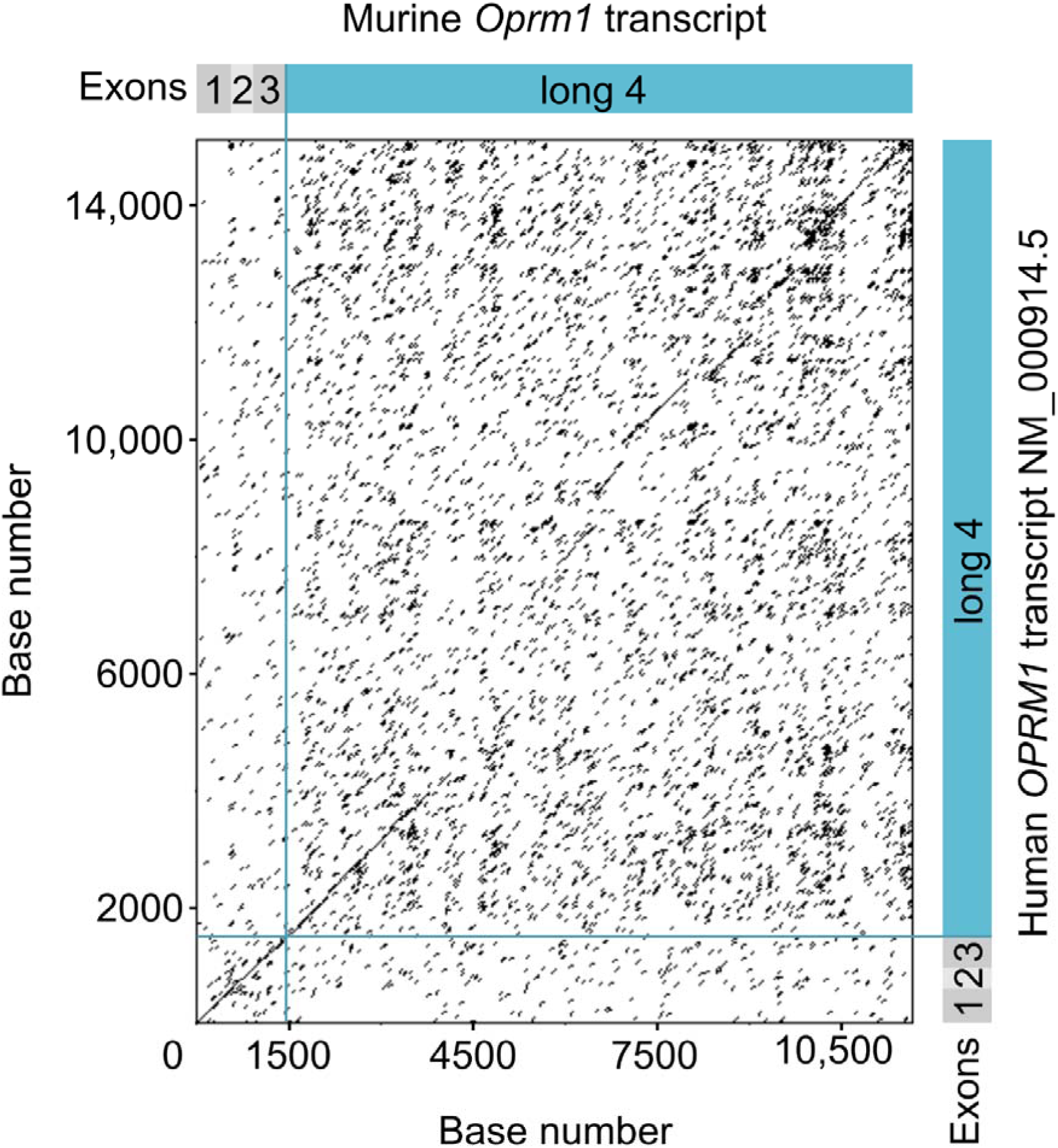
Similarity between human and murine variants of *Oprm1* transcripts with long exon 4. The dot plot illustrates the similarity of the primary human transcript variant ENST00000330432.12 and the primary murine transcript observed here. The comparison revealed regions of high conservation within the first 3500 bp of the mouse transcript. The region contains a 5’UTR upstream of exon 1, a protein-coding sequence, and an ~2 kb long region of the 3’UTR. Additionally, several highly conserved regions in the distal 4 kb of the mouse 3’UTR were detected. Analysis was performed using EMBOSS/dotmatcher (www.ebi.ac.uk/tools/emboss), with a window size of 45 nucleotides and a threshold of 35.

### Cellular distribution of the long 30 UTR in the selected brain regions and spinal cord

To confirm the presence and relative abundance of the long *Oprm 1* variant, we used fluorescent *in situ* hybridization with a set of variant-specific RNAscope probes. Three probes were used, the first targeting essential exons 2 and 3, the second targeting the distal part of the long 3 UTR (chr10:6,858,777-6,860,133), and the third targeting a 3 terminal region including exons 7, 19, 8, and 9. Analyses were performed on coronal sections of the brain, including the nucleus accumbens, striatum, prefrontal cortex, primary motor cortex, primary somatosensory cortex and coronal sections of the spine at the L3-5 level. As illustrated in **Figures 5 and 6**, multiple signal puncta in the perinuclear region were detected with the first and second probes across all analyzed areas. The staining was primarily localized in the proximity of the DAPI-stained nuclei. Conversely, relatively few signal puncta were observed for the third probe, and the signal was scattered. In the spinal cord, signals from probes against exons 2 and 3 or the long exon 4 were abundantly present in the dorsal and ventral horns, in line with the known distribution of *Oprm1* expression (Mansour, Fox, et al., 1995). The signal from the probe targeting exons 7, 19, 8, and 9 was appreciably lower than that from the forebrain. Multiple overlapping puncta were detected with probes targeting exons 2 and 3 and long exon 4 in all areas analyzed. Quantitative analysis of the images revealed the greatest number of *Oprm1* transcripts in the nucleus accumbens and striatum, a relatively high number in the spinal cord, and the lowest number in the cortical areas (**Figure 7**). The greatest number of puncta was detected with the second probe against long exon 4, and notably, in the spinal cord, it was more than twice that observed for the probe against exons 2 and 3. The signals from probes targeting exons 7, 19, 8, and 9 in the spinal cord were similar to those in the negative controls (**Figures S2&S3**). The overlap between probes against exons 2 and 3 and against long exon 4 was between 44.6% and 57% in all brain and spinal cord areas analyzed. Conversely, the greatest overlap between probes targeting exons 2&3 and exons 7, 19, 8, and 9 was in cortical areas—up to 51% in the somatosensory cortex, 12% and 13% in the dorsal striatum and nucleus accumbens, respectively—and approximately 6% in the spinal cord. We also detected overlap between probes targeting long exon 4 and exons 7, 19, 8, and 9, which had the same trend in distribution as in the previous case, i.e., highest in the cortex (up to 40%), followed by the striatum and nucleus accumbens (23% and 18%, respectively), and lowest in the spinal cord (~3%). The overlap between probes targeting long exons 4 and exons 7, 19, 8, and 9 was unexpected, as the existence of a transcript including long exon 4 and some or all of exons 7, 19, 8, and 9 was not supported by the sequencing data. Notably, when signal colocalization in the positive and negative controls was analyzed (**Figures S2&S3**), the colocalization was relatively low in the case of abundant positive controls and high among negative controls, with a minimal number of detected puncta. Thus, a lower signal intensity may be associated with greater nonspecific colocalization. Therefore, we argue that the data clearly show colocalization on the same transcript of probes against exons 2 and 3 and long exon 4. However, the interpretation is less clear in the case of probes targeting exons 7, 19, 8, and 9. Sequencing data support the existence of isoforms containing some or all of these exons, and thus, the degree of colocalization probably reflects the presence of alternative transcripts; nevertheless, based on negative controls, we assume that a fraction of this colocalization may be artifactual.

**Figure 5.**
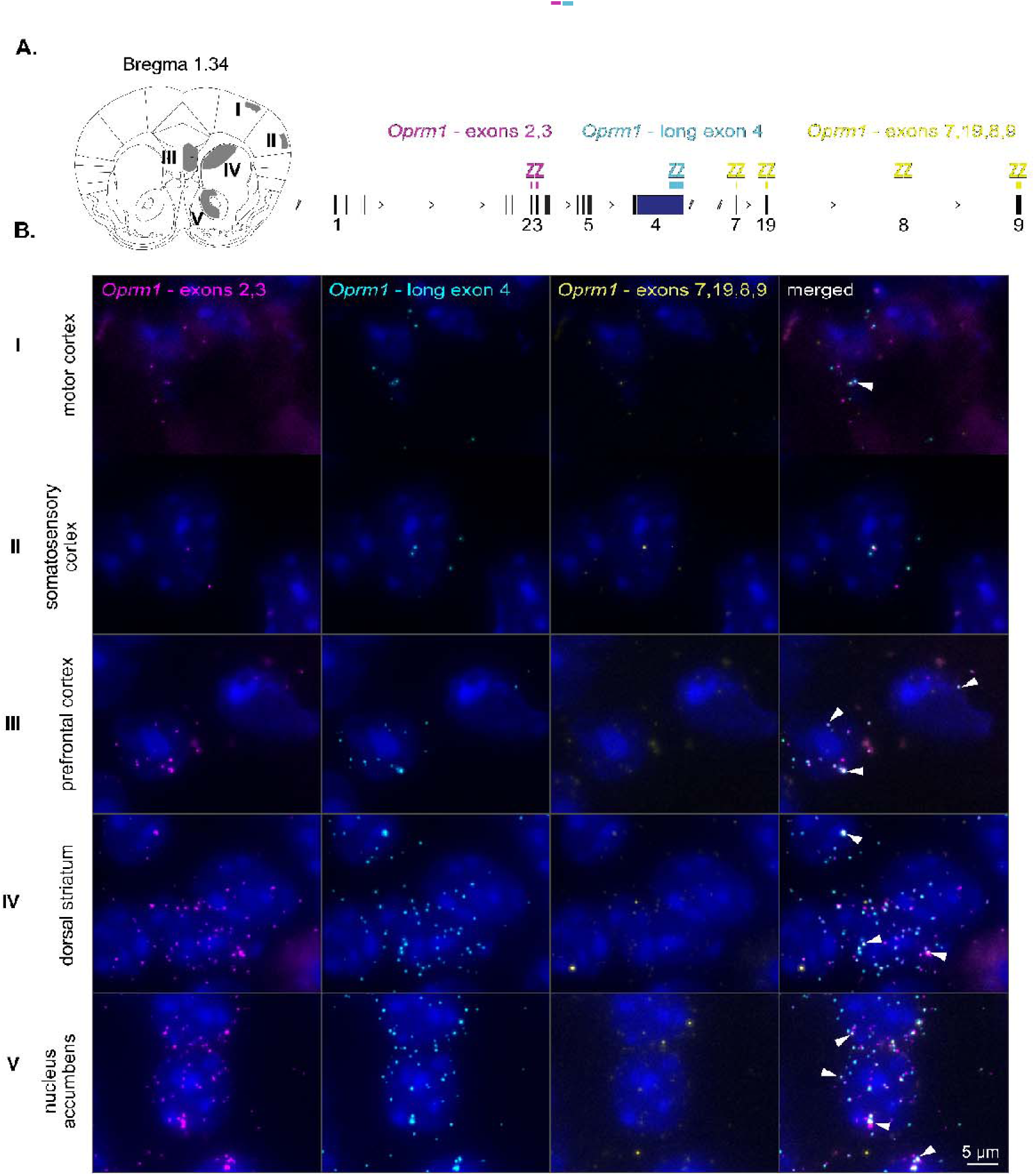
*In situ* analysis of *Oprm1* transcripts in the mouse forebrain. **A.** A schematic representation of the brain areas analyzed by RNAscope *in situ* hybridization (left) and *Oprm1* regions targeted by the 3 probes used (right). **B.** Representative micrographs for each of the examined brain structures. Maximum intensity projections for separate channels, as well as merged photos, are presented. Yellow arrows indicate *Oprm1*-exons 2, 3 and *Oprm1*-long exon 4 colocalized particles, white arrows indicate *Oprm1*-exons 1,2 and *Oprm1*-exons 7-9 colocalized particles. The scale bar is 5 µm.

**Figure 6.**
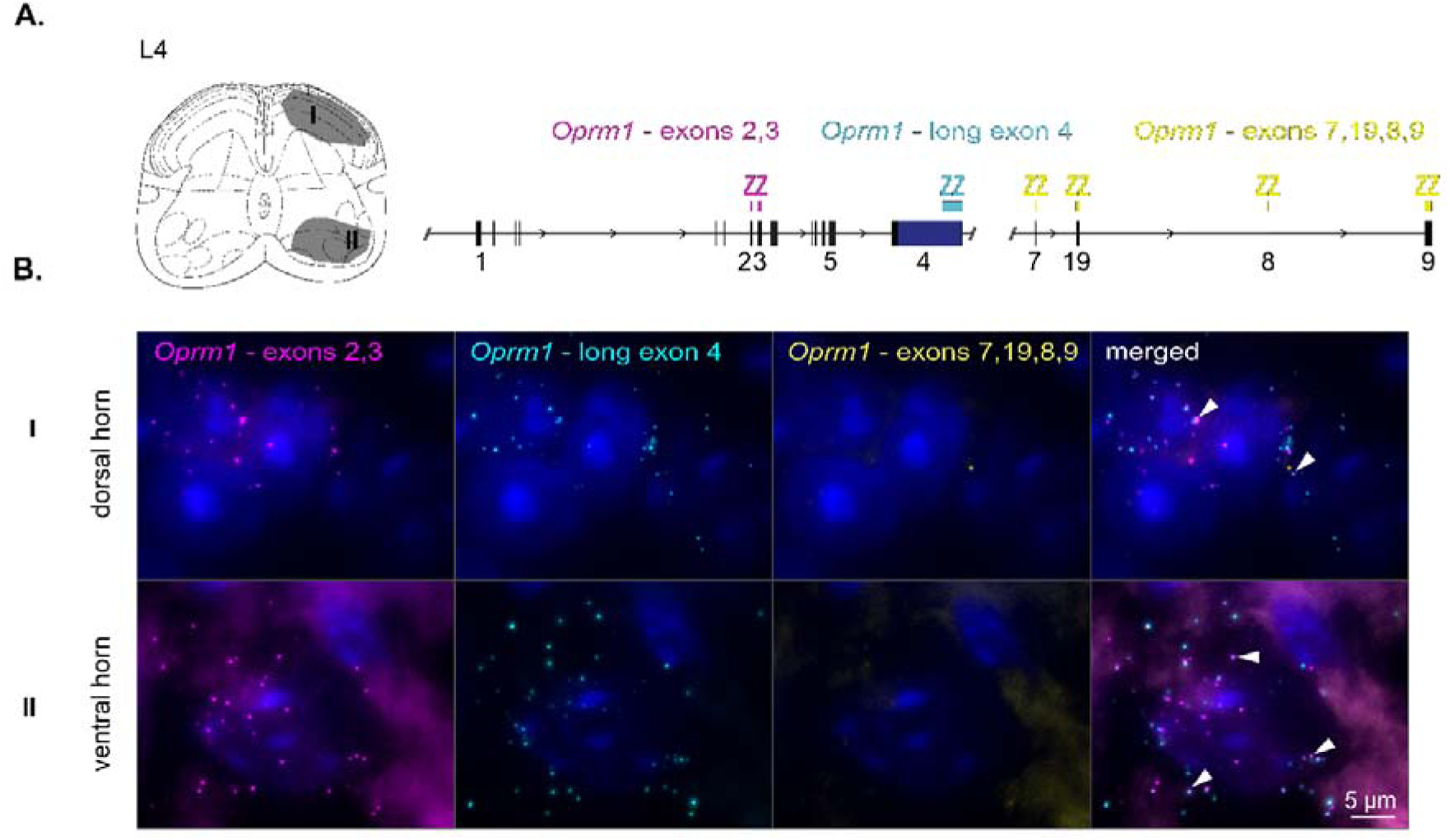
*In situ* analysis of *Oprm1* transcripts in the spinal cord. **A.** A schematic representation of the spinal cord areas analyzed by RNAscope *in situ* hybridization (left) and *Oprm1* regions targeted by the 3 probes used (right). **B.** Representative micrographs of the dorsal and ventral horns at the L4 level. Maximum intensity projections for separate channels, as well as merged photos, are presented. Yellow arrows indicate *Oprm1*-exons2,3 and *Oprm1*-long exon 4 colocalized particles, white arrows indicate *Oprm1*-exons1,2 and *Oprm1*-exons7-9 colocalized particles. The scale bar is 5 µm.

**Figure 7.**
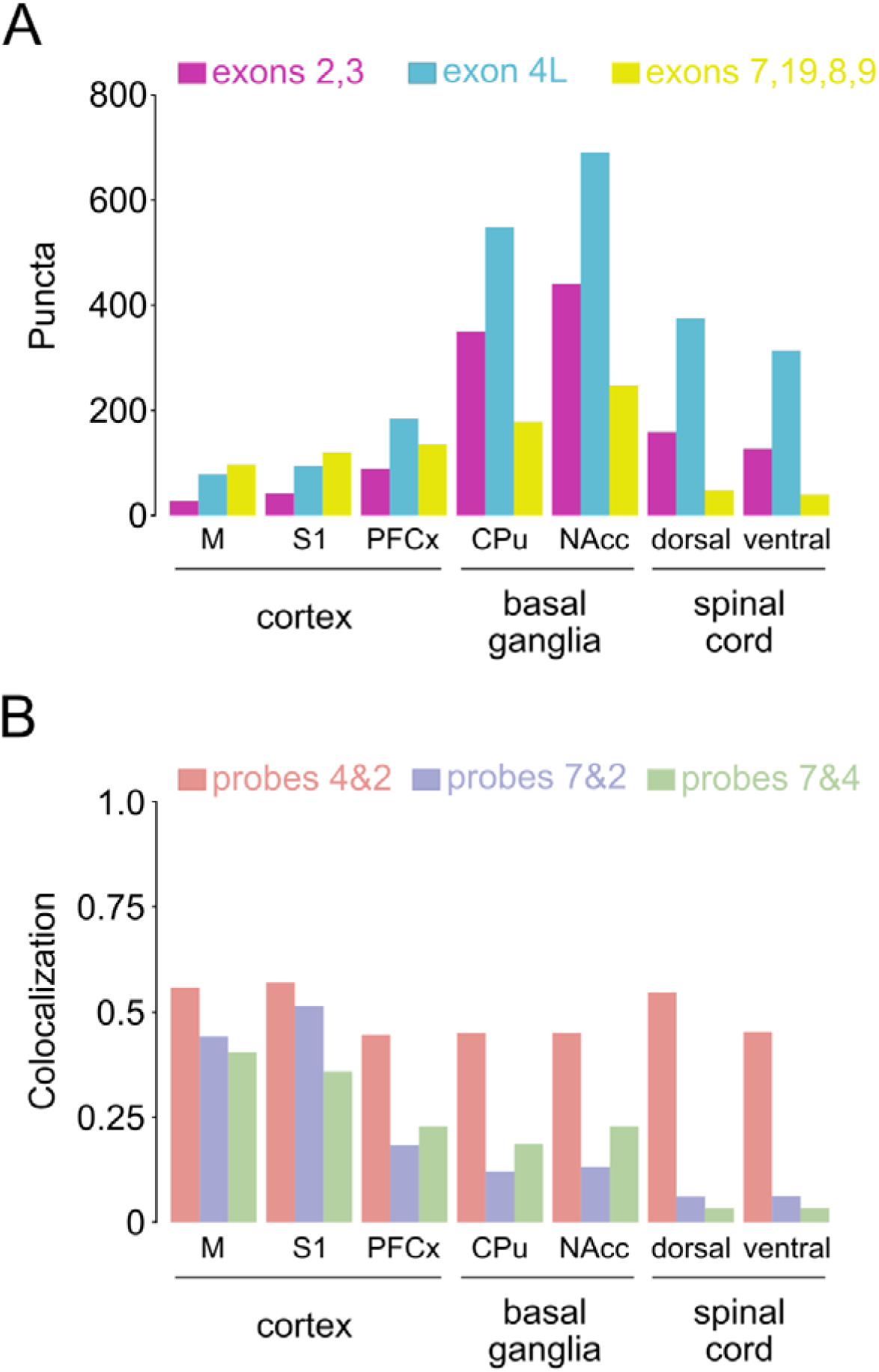
Colocalization of probes targeting different regions of *Oprm1* transcripts. The panel at the top shows exons targeted by the 3 RNAscope probes. A. Summary of the number of mRNA puncta. The bars show the number of puncta counted for each probe in 4 separate micrographs corresponding to the areas indicated below the graph, and the colors represent probes as indicated above. B. Overlap of puncta detected with different RNAscope probes. Bars represent fractions of the probe with fewer puncta that colocalized with the probe with a larger puncta count, i.e., the first probe to the second probe indicated in the color legend. Abbreviations: M – motor, S1 – primary somatosensory cortex, PFCx – prefrontal cortex, CPu – caudate and putamen, NAcc – nucleus accumbens septi.

### Expression of Oprm1 in different cortical cell types

To independently validate the results of the analysis and possibly determine the cell types expressing the *Oprm1* gene, we reanalyzed single-cell transcriptomic data from Allen’s V1 & ALM - SMART-SEQ dataset (Tasic et al., 2018). The sets included cells extracted from the primary visual cortex (VISp) and the anterior lateral motor cortex (ALM). Analysis revealed that the *Oprm1* gene was expressed mainly in parvalbumin-positive, somatostatin-positive, and Vip-positive cells. Relative expression of *Oprm1* exons 1, 2, and 3 greater than 0.01 was also detected in subsets of cortical layer 5 and 6 neurons. Sequence reads aligned to exons 1, 2, and 3 and the long 4 variant were detected in all GABAergic neurons, with examples of specific subtypes shown in **Figure 8**. Reads corresponding to canonical short exon 4 and exon 5 were also observed, as well as a low frequency of reads aligned to exons 10a, 15d and 19. Thus, there is clear evidence of a primary *Oprm1* transcript comprising exons 1, 2, and 3 and a long variant of 4. Additionally, low levels of reads of exon 19 agreed with both sequencing and RNAscope data. Taken together, the results of all the analyses performed show that the main transcript of the *Oprm1* gene in mice contains a very long 3 terminal exon, with limited evidence for alternative 3 variants and no alternative 51 variants.

**Figure 8.**
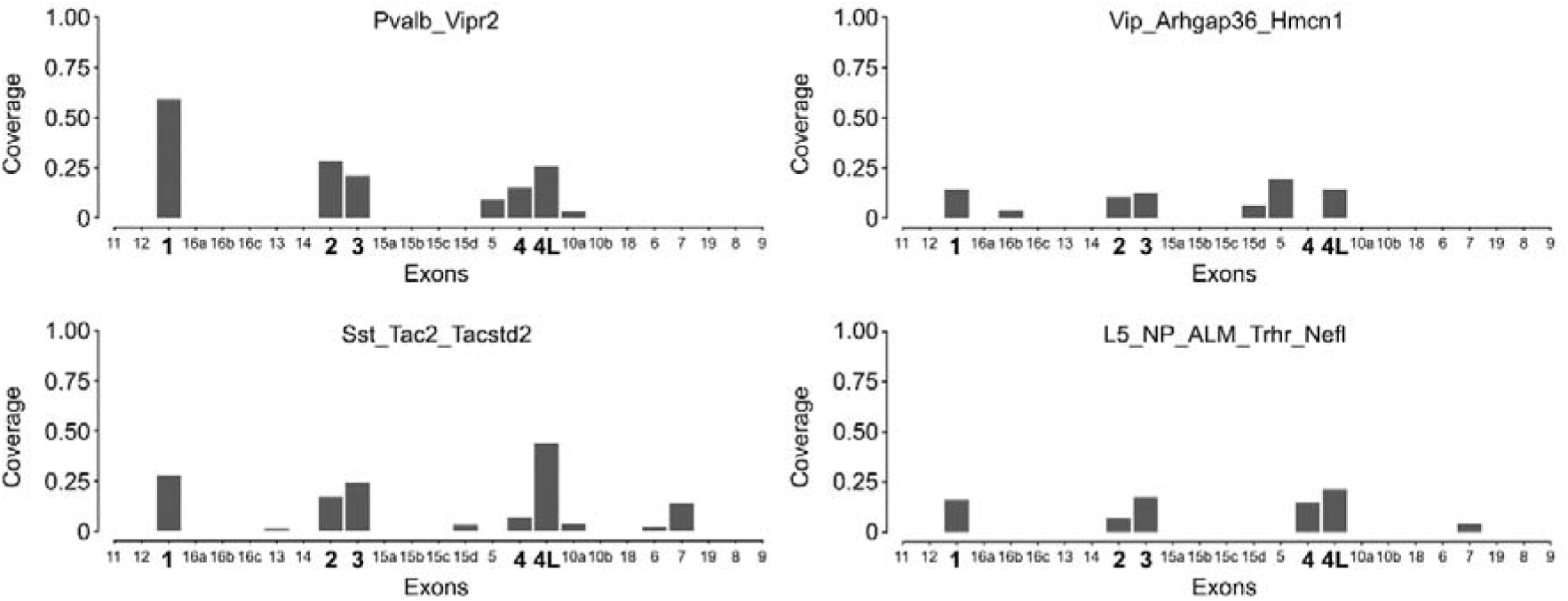
Alignment of sequence reads from the Allen Atlas cortical single-cell to putative *Oprm1* exons (Tasic et al., 2018). The diagram at the top represents reported exons and long exon 4. The graphs show the relative coverage of *Oprm1* exons in 4 types of cells selected for high coverage of exons 1 to 3. The names above the graphs represent cell subtypes in the Allen Atlas classification. Pvalb-, Vip-, and Sst-represent parvalbumin-, vasoactive intestinal polypeptide (VIP)- and somatostatin-expressing GABAergic cells, respectively. L5 cells are a subtype of glutamatergic layer 5 cortical neurons. Exons 1 to 4 and the long variant of exon 4 have larger labels on the X-axis.

## Discussion

We found that the primary transcript of the murine *Oprm1* gene has a long 3 terminal exon, which is homologous to the recently revised human *OPRM1* reference transcript (NM_000914.5). The presence of the long transcript was validated through long-read sequencing, 3 termini sequencing, reanalysis of public datasets, and *in situ* hybridization. The observed long 3 terminal exon in the primary *Oprm1* transcript is consistent with previously reported observations (Ide et al., 2005; Ikeda et al., 2001; Wu et al., 2005). Compared to previously reported data, there are minor differences in the sequence of the long 3 terminus, and we observed slightly greater homology to the human reference transcript. In all the analyses performed, the long *Oprm1* variant was the predominant transcript, accounting for at least 85% of the total transcription. Evidence supporting the existence of alternative transcripts that include 3’ terminal exon 9 was also observed. Conversely, long-read sequencing revealed no evidence of variants with alternative transcription starts and no variants affecting the region encoding the transmembrane helices (i.e., exons 1, 2 and 3). The long 3 terminal exon described here encodes 12 amino acids at the C-terminus of the μ-opioid receptor (‘LENLEAETAPLP’), and this sequence is identical to the C-terminus encoded by the human reference *OPRM1* transcript (Liu et al., 2021).

There are two important implications of our results for further research on opioid receptor ligands. First, we confirmed that the primary mouse and human opioid receptor μ transcripts are homologous and may be involved in similar molecular regulatory mechanisms. Therefore, the effects of opioids observed in mice should have robust predictive value with regard to potential actions in humans. Our results support the existence of a small fraction of *Oprm1* variants with alternative C-termini, although their levels were close to the detection limit in all analyses, and we argue that the data show no discrete pattern in their distribution. The *in sit u*analysis suggested that the alternative variants are more abundant in the cortex; nevertheless, we remain cautious, as the result resembled the colocalization observed in the case of negative controls. Thus, our second conclusion is that no evidence of transcripts encoding alternative sequences involved in ligand binding or G-protein or β-arrestin interactions was found, which limits the feasibility of developing novel opioids targeting specific receptor subpopulations. Moreover, these data suggest that different protein variants are unlikely to account for previously reported pharmacologically different opioid receptor populations (Narayan et al., 2021; Pasternak & Pan, 2013; Xu et al., 2017).

The function of the long 3 terminal transcript sequence described here remains elusive. Previous reports indicated a role for the long *Oprm1-*3’UTR in opioid sensitivity (Ikeda et al., 2001) and assessed the presence of repeated sequence motifs. In recent years, the transcript sequences of thousands of genes in both rodents and humans have been updated with longer 3 UTRs (Miura et al., 2013). Substantially longer 3 UTRs were observed in neurons. Many of these transcripts were shown to possess >10 kb 3 UTRs, and some of them exceeded >18 kb. It has been suggested that this region contains sequences responsible for the regulation of gene expression, stability and cellular localization of mRNAs (Bae & Miura, 2020). For instance, analysis of the rodent hippocampus showed that the 3 UTRs found in the neuropil influenced longer mRNA half-lives than did the 3 UTRs in the somatic compartment (Tushev et al., 2018). Conversely, knockout of this region in the mTOR gene significantly limits the distribution of mRNAs to specific neuronal compartments (Terenzio et al., 2018). The *in situ* hybridization analysis presented here showed that *Oprm1* transcripts clustered in the perinuclear area; however, to properly assess the distribution of transcripts, it would be necessary to visualize cell processes and possibly dendritic spines.

Taken together, our data show that the *Oprm1* gene has one primary transcript with a very long 3 untranslated sequence. All four exons of the primary transcript are homologous to the reference transcript of human *OPRM1*. Low levels of a secondary transcript with an alternative 3 terminus were detected; however, its abundance was low, and it was not consistently detected by all methods employed. The function of the long 3 terminal region remains unknown, although its presence appears to be consistent with that of several other neuronal transcripts.

## Methods

### Oprm1 transcript comparison

The data used for transcript comparison (**Figure 1**, **Table S1**) were obtained from Ensembl (Ensembl release 108 - Oct 2022) and the National Center for Biotechnology Information (NCBI) GenBank Release 253: December 15 2022. For the NCBI search, the following search terms were used: ‘(mu opioid receptor[Title]) AND *Mus musculus* [Organism]’, and ‘(Oprm1[Gene Name]) AND *Mus musculus* [Organism]’. The resulting sequences were then aligned to the GRCm38/mm10 mouse genome using the BLAT tool (Kent, 2002). **Figure 1** was created using the ggtranscript extension for ggplot2 (https://github.com/dzhang32/ggtranscript) and R version 4.2.2. The output was graphically modified using Inkscape v1.2.2.

### Single-cell dataset reanalysis

Reanalysis of single-cell RNA-seq datasets from Tabula Muris (Schaum et al., 2018) and Allen’s V1 & ALM - SMART-SEQ (2018) experiments was performed using the ‘bigWigSummary’ script from the UCSC Genome Browser. The script was used to calculate the average values of genome coverage in the regions that were previously identified as *Oprm1* exons. In case of exons with multiple start/stop positions, the coverage was assessed for the narrowest exon segments (Supplementary Materials, Table 1).

### Oxford Nanopore sequencing

Tissue samples were collected from C57BL/6 mice. Experimental mice were housed at the animal facility of the Maj Institute of Pharmacology of the Polish Academy of Sciences and were euthanized between 10–18 weeks of age by cervical dislocation. Both male and female animals were included in the long-read analysis (ONS), and only males were used for spatial transcriptomics and *in situ* hybridization.

The mouse brains were removed, immersed in RNAlater Stabilization Solution (Qiagen Inc., USA) and stored at 4 °C. The next day, slices containing the striatum and prefrontal cortex (anterior to bregma 0.74 to 1.34, based on Paxinos & Franklin, 2001), as well as the thalamus (posterior to bregma −0.94 to −1.82, based on Paxinos & Franklin, 2001), were cut on a vibratome VT1200 (Leica, Germany) into 200 μm sections. Additionally, the thalami were dissected from the slices under a stereomicroscope (StemiDV4, Carl Zeiss, Germany) using sterile needles. The tissue samples were placed in 2.0 ml round bottom Eppendorf® tubes and kept at −20 °C.

RNA was isolated using the single-step acid guanidinium thiocyanate-phenol-chloroform extraction method (Chomczynski, 1993). The samples were thawed at room temperature, the RNAlater solution was removed, and stainless steel balls were placed into the tubes. First, tissue samples were homogenized in 1 ml of TRIzol reagent (Invitrogen, USA) in a TissueLyser II apparatus (Qiagen Inc., USA; 2 × 3 min, 25 Hz). To remove phenol traces, 200 µl of chloroform was added to each tube. The samples were incubated for 10 minutes on ice, mixed by vortexing, and centrifuged (12 000 × g, 20 minutes, 4 °C). The aqueous phase (approx. 450 µl) was transferred to new 1.5 ml tubes, and an equal volume of isopropanol (POCH, Poland) was added to each tube. The samples were mixed, placed at −70 °C for 20 minutes and then centrifuged (12 000 × g, 30 minutes, 4 °C). The supernatant was discarded, and the RNA-containing pellets were resuspended in 1 ml of 70% (v/v) ethanol and further centrifuged (12 000 g, 10 minutes, 4 °C). After washing with ethanol, the pellets were air-dried and dissolved in 30–50 µl of sterile, nuclease-free water (Ambion Inc., USA). Finally, the samples were incubated in a thermomixer (Thermomixer Comfort, Eppendorf, Germany) set at 65 °C for 5 minutes and then inverted several times. After RNA isolation, RNA integrity was determined using capillary chip-based electrophoresis with an RNA 6000 Nano Methods 59 LabChip Kit and an Agilent Bioanalyzer 2100 (Agilent, USA) according to the manufacturer’s instructions.

Reverse transcription was performed using the Omniscript RT Kit (Qiagen Inc., USA). RNA samples were thawed on ice, mixed by vortexing and briefly spun down at 4 °C. Then, 10 × Buffer RT, dNTP mix, RNase-free water and oligo-dT primers (Invitrogen, USA) were thawed at RT, mixed, short spin centrifuged, and kept on ice. The components were added according to the manufacturer’s instructions to obtain 2 µg of template RNA in 20 µl of the reaction mixture. The 10 × Buffer RT, dNTP mix, oligo-dT primers and Omniscript Reverse Transcriptase enzyme were mixed together, added to tubes with diluted template RNA, and gently stirred with a pipette. The samples were incubated for 60 minutes at 37 °C and then diluted 20 times with sterile water. The obtained cDNA samples were subsequently sent for sequencing.

### Sequencing analysis

Sequencing was performed by Novogene UK using the Nanopore PromethION platform. Each sample was sequenced three times. The raw statistics that describe the quality and characteristics of the procedures are presented in **Table S2**. The number of sample bases varied between 5.2 and 11.5 Gbp per sample, while the number of reads fluctuated between 3.8 and 8.2 × 10^6^. The median read length in all the samples was relatively consistent and ranged between 1 002 and 1 165 bp. The N50 read length, which is the length of the shortest read in the subset of the longest sequences that together represent ≥50% of the nucleotides in the sample, varied from 1 875 to 2 172 bp, with greater values obtained in the samples containing the striatum and prefrontal cortex than in those containing the thalamus (**Table S2**). The median read quality was high and consistent between samples, reaching values between 12.1 and 13.0. The percentage of reads with relatively low quality (<7) did not exceed 3.5% in any of the samples. Sequencing data quality was checked with MinIONQC v1.4.2 and R version v4.1.3. All FASTQ files for each brain region were merged, and the data were aligned to the GRCm38_102/mm10 genome with the Minimap2 program (v. 2.24). The alignment results were filtered for strand mismatches and invalid 3’ ends. The final number of valid alignments was ~35 million. The output files were visualized using Integrative Genome Viewer v2.12.3 (IGV; Thorvaldsdóttir et al., 2013). Raw sequence reads were deposited in the Sequence Read Archive: https://www.ncbi.nlm.nih.gov/bioproject/PRJNA1032908.

### Spatial transcriptomics dataset

The dataset was generated as part of a separate project focused on the effects of L-DOPA on gene expression in mice with progressive loss of dopaminergic neurons (Radlicka-Borysewska, in preparation) and is available from the Sequence Read Archive database: https://www.ncbi.nlm.nih.gov/bioproject/PRJNA1080215. The procedure was performed following the protocol from the Visium Spatial Gene Expression User Guide Revision E (10x Genomics, USA). Briefly, mouse brains were dissected, embedded in OCT medium (CellPath, United Kingdom), and flash frozen using isopentane and liquid nitrogen. The brains were stored at −80°C. On the day of the experiment, 10 μm thick slices from the rostral part of the forebrain (Bregma 1.18 to 1.98 mm, based on Paxinos & Franklin, 2001) were obtained using the CM 3050 S cryostat (Leica, Germany). 12 sections were mounted on Visium slides (10x Genomics, USA) containing spots with poly(dT) primers. Forebrain sections were first fixed, stained with hematoxylin and eosin, and imaged separately via bright field microscopy using a Leica DMi8 system (Leica, Germany). After imaging, the slides were processed for cDNA synthesis and amplification following the manufacturer’s instructions. First, the tissue was permeabilized, allowing the mRNA to hybridize with poly(dT) primers. Subsequently, cDNA was synthesized, the second strand was generated, and then the strands were denatured. A quantitative polymerase chain reaction (qPCR) was conducted on a portion of each sample to assess the number of cycles sufficient for amplification. After amplification was performed, cDNA was purified and used for library construction. Libraries were checked for quality and quantity and sent for sequencing on a NovaSeq 6000 instrument (Illumina, USA). Raw results were analyzed using Space Ranger v1.3.1, and subsequently MACS3 (https://github.com/macs3-project/MACS, Zhang et al., 2008). The method allowed for the alignment of RNA-seq reads to specific genes or long terminal repeats described in the GRCm38/mm10 genome. As a result, ‘peaks’ of reads were annotated to the genome. The method also provided information on the DNA strands from which the annotated RNA-seq reads originated. The coordinates of the obtained peaks were compared with the data obtained in the Oxford Nanopore Sequencing experiment.

### RNAscope *in situ* hybridization

RNAscope (Advanced Cell Diagnostics, Inc., ACD) with three probes, Mm-Oprm1-O7-C1, Cat No. 1178981-C1, Mm-Oprm1-O5-C2, Cat No. 568771-C2, and Mm-Oprm1-O4-C3, Cat No. 544731-C3, was used to detect *Oprm1* exon-specific expression in different murine brain regions. The animals were killed by cervical dislocation. The brains were removed immediately from the skulls, fresh frozen on dry ice, embedded in optimal cutting temperature (OCT) compound (Cell Path, UK), and stored at −80 °C for up to 2 months. Frozen brains were sliced into 10 µm coronal sections on a cryostat (CM 3050 S, Leica, Germany) with both the object and chamber temperature set at −20 °C. The slices were thaw-mounted on positively charged SuperFrost Plus microscope slides and stored at −80 °C for up to one month. An RNAscope fluorescent multiplex assay was performed according to the manufacturer’s instructions. Slices representing the dorsal striatum, nucleus accumbens, prefrontal cortex (the cingulate cortex and infralimbic cortex), primary motor cortex and primary somatosensory cortex as well as the lumbar spinal cord were selected for ISH. The RNAscope assay started with prefixing the slices with ice-cold 4% paraformaldehyde (PFA), followed by dehydration at increasing concentrations of ethanol. After dehydration, the slides were air-dried, and a hydrophobic barrier was drawn. To permeabilize the tissue, protease IV was applied for 30 minutes at room temperature. The slides were then washed in 1× PBS and hybridized with a specific probe for 2 h at 40 °C. After this step, four hybridizations with subsequent amplifiers were performed. In the final hybridization step, fluorophores were attached to the targets. After that, the specimens were counterstained with DAPI. The coverslips were mounted with ProLong Gold Antifade Mountant (Invitrogen, USA), and the slides were stored at 4 °C in an opaque box.

Image acquisition was performed on the day following the ISH procedure. For imaging, an Axio Imager.Z2 fluorescence microscope (Carl Zeiss, Germany) with Plan-Apochromat 63×/1.40 Oil M27 lenses and Axiocam 506 camera was used. Laser lengths equal to 644 (At647), 553 (At550), 493 (AF488), and 353 nm (DAPI) were used to excite the fluorophores. The laser power was adjusted to the brightest sample and remained the same throughout the experiment. For each stage, 21 5 μm thick Z-stacks were acquired.

Image processing was conducted using ZEN Lite Software (version 3.5.093.00002, Carl Zeiss, Germany). For each Z-stack, the maximum orthogonal projection was calculated using Zeiss ZEN Software. Images that were out of focus were excluded from the input file. The maximum projection images were then analyzed with the Fiji/ImageJ plugin ComDet v.0.5.5 (Katrukha, 2020). At647, At550, and Af488 particles per channel were detected in the regions of interest based on their size and intensity threshold compared to the image background. Particle colocalization was calculated based on the maximum distance between the particles.

---

**Figure S1.**
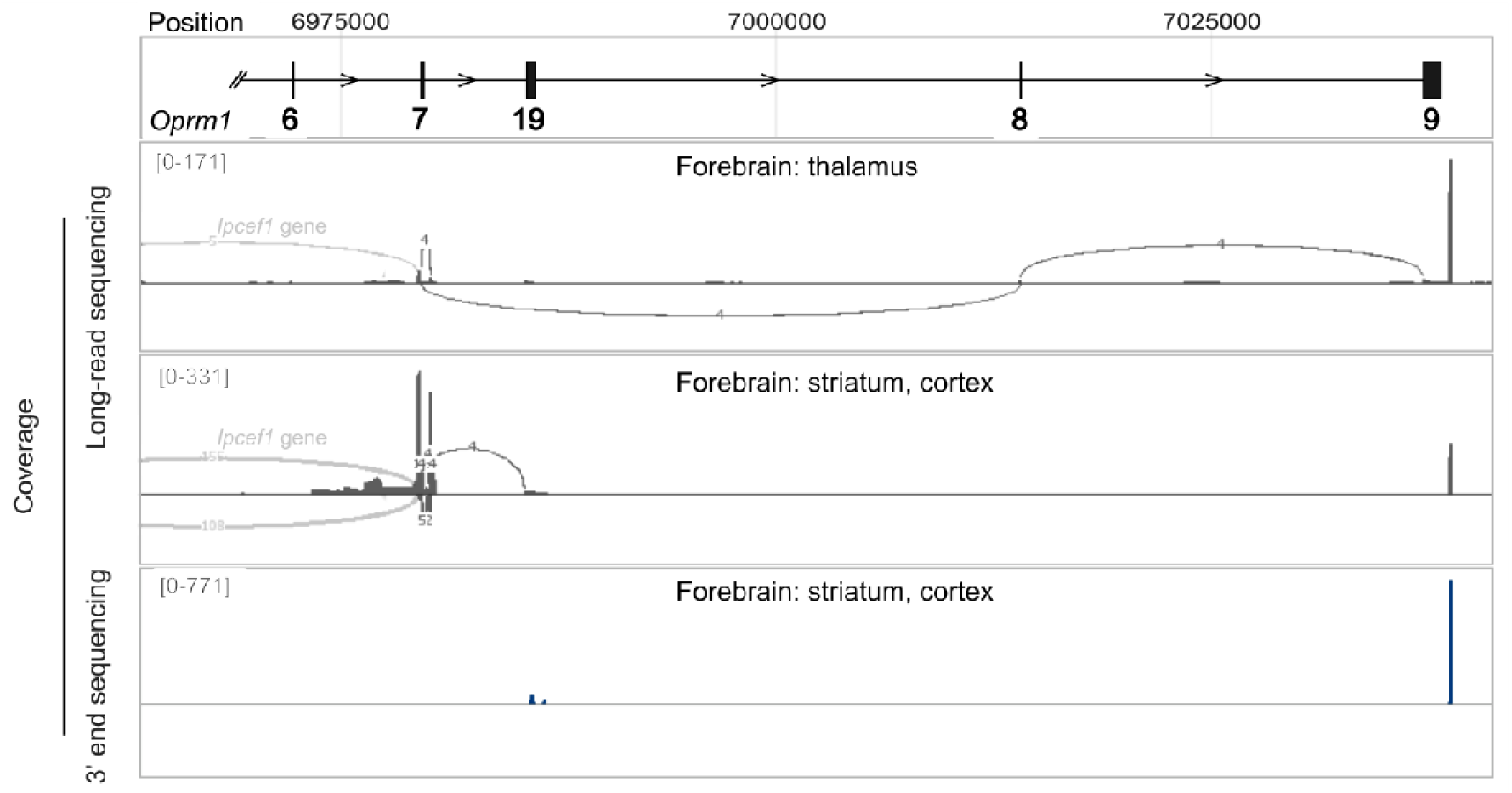
Alternative transcripts of the *Oprm1* gene. The diagram at the top shows the positions of putative exons in the 3’ region of the gene. The first two lanes show “Sashimi” plots of long-read sequencing analysis of brain regions, including the cortex and striatum (the first lane) and the thalamus (the second lane). The third lane shows sequencing results based on spatial RNA sequencing of 3’ termini in a frontal brain coronal section. Reads corresponding to the region expanding beyond exon 4 and their position in the schematic sequence at the bottom are shown in magenta.

**Figure S2.**
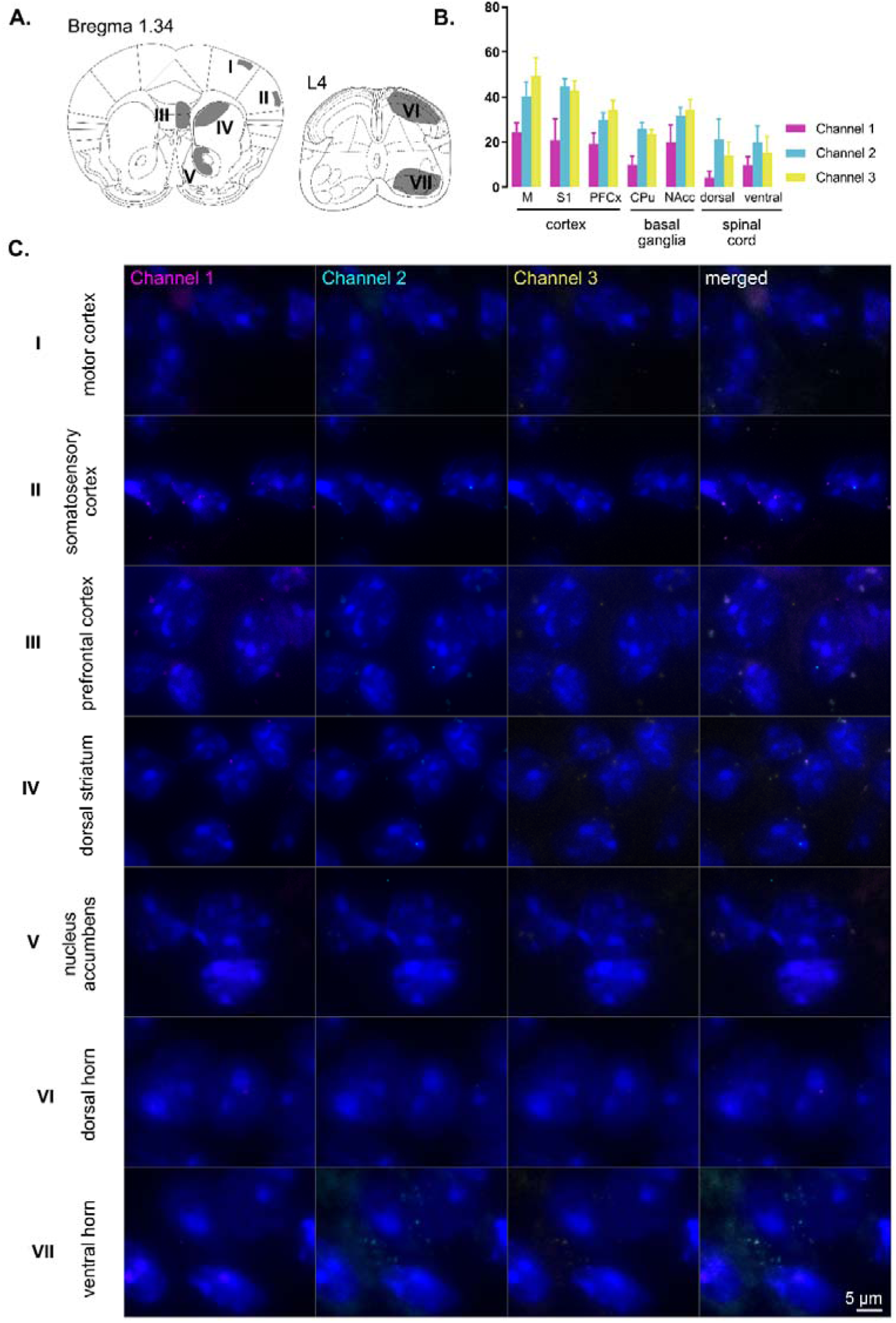
Negative controls for in situ RNAscope hybridization. **A.** A schematic representation of the brain and spinal cord areas analyzed. **B**. The number of mRNA puncta counted for each of the negative control probes. **C.** Representative micrographs for each of the examined brain structures. Maximum intensity projections for separate channels, as well as merged photos, are presented. The scale bar is 10 µm. Abbreviations: M – motor, S1 – primary somatosensory cortex, PFCx – prefrontal cortex, CPu – caudate and putamen, NAcc – nucleus accumbens septi.

**Figure S3.**
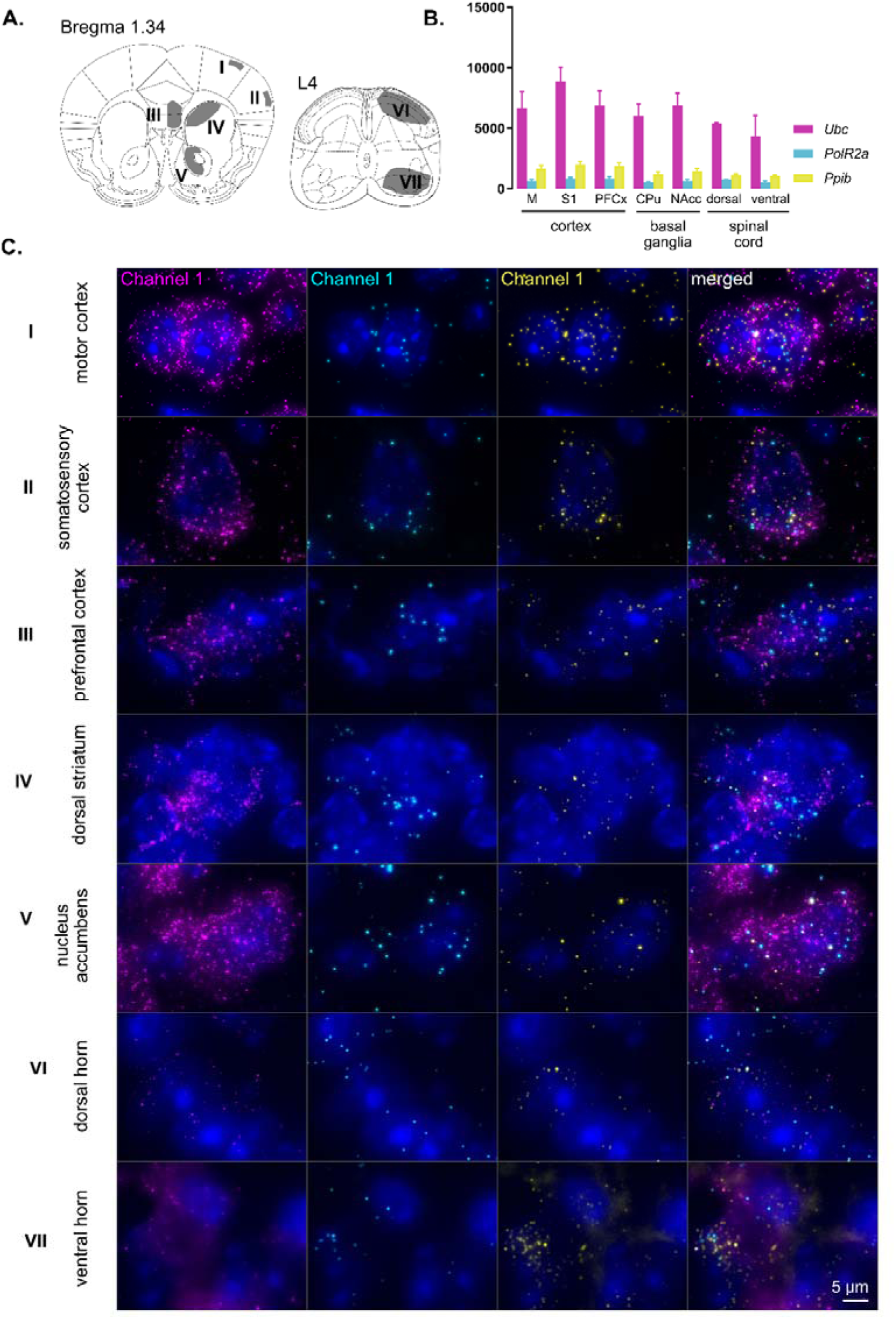
Positive controls for in situ RNAscope hybridization. **A.** A schematic representation of the brain and spinal cord areas analyzed. **B**. The number of mRNA puncta counted for each of the positive control probes. **C.** Representative micrographs for each of the examined brain structures. Maximum intensity projections for separate channels, as well as merged photos, are presented. The scale bar is 10 µm. Abbreviations: M – motor, S1 – primary somatosensory cortex, PFCx – prefrontal cortex, CPu – caudate and putamen, NAcc – nucleus accumbens septi.

## Notes

### Competing Interest Statement

The authors have declared no competing interest.

